# Reproductive skew affects social information use

**DOI:** 10.1101/410886

**Authors:** Marco Smolla, Charlotte Rosher, R. Tucker Gilman, Susanne Shultz

**Affiliations:** Department of Biology, University of Pennsylvania, Philadelphia, PA, United States of America; School of Earth and Environmental Sciences, Faculty of Science and Engineering, University of Manchester, United Kingdom; Department of Ecology & Genetics, Uppsala University, Sweden

**Keywords:** cultural evolution, reproductive skew, social learning, competition, agent-based model

## Abstract

Individuals vary in their propensity to use social learning, the engine of cultural evolution, to acquire information about their environment. The causes of those differences, however, remain largely unclear. Individuals that experience high reproductive skew are expected to favour high-risk strategies, whereas those that experience low reproductive skew are expected to favour risk-averse strategies. Using an agent-based model, we tested the hypothesis that differences in energetic requirements for reproduction affect the value of social information. We found that social learning is associated with lower variance in yield and is more likely to evolve in risk-averse low-skew populations than in high-skew populations. Reproductive skew may also result in sex differences in social information use, as females tend to be more risk averse than males. To explore how risk may affect sex differences in learning strategies, we simulated learning in sexually reproducing populations. Where both sexes share the same environment they adopt more extreme learning strategies, approaching pure individual or social learning. These results provide insight into the conditions that promote individual and species level variation in social learning and so may affect cultural evolution.

## 1 Introduction

Social learning plays a central role in the emergence of traditions and the evolution of culture, as it allows locally adaptive information and behavioural traits to spread across a population [1]. How well information is transmitted throughout a population depends on the fidelity and frequency of social learning [2]. If only a small subset of a population learns socially or social learning occurs infrequently, information is expected to spread more slowly, have a higher probability of being lost, and perhaps be used less efficiently [2, 3]. Making useful predictions for how information emerges and diffuses in social groups, therefore, relies on a good understanding of social learning dynamics and how it varies across a population [4].

One source of variation in social learning that has been intensely studied in the past is that socially acquired information is not universally beneficial [4-6]. While social information can be acquired faster or at a lower cost, it is also prone to be outdated, less relevant or less reliable than individually acquired information [7, 8]. Accordingly, social learning is often used in a context-dependent manner, following a range of behavioural rules that describe when to learn socially and when to learn individually [9, 10]. These social learning strategies have been found across the animal kingdom in a variety of contexts, including the acquisition of food, mate and nest-site choice preferences, foraging techniques and predator avoidance behaviours [7, 8, 11].

Beyond context-dependency, another source of variation in social learning is inter-individual differences, such as personality traits, reproductive state, and developmental stress, as well as age, sex, and dominance [12-17], which have been reported for a variety of species [18]. Previous empirical work provided different explanations for the observed inter-individual patterns of social learning. For example, a study on meerkats found individual differences in social learning based on age [15]. Young meerkats acquired social information more readily than older individuals. The authors suggest that this is not due to differences in learning capacity, but rather because young meerkats rely more strongly on food and protection provided by adults, and so have more opportunities to observe and learn from adults. A study on guppies found differences based on sex, where foraging information spread faster in groups of females than in groups of males [16]. This was likely due to female reproductive success being mostly limited by access to resources, whereas male reproductive success was limited by access to females. Another study on fish found differences based on reproductive state [13]. Gravid female sticklebacks relied more strongly on social information to find a rich food patch than did males or non-gravid females, whereas reproductive males disregarded social information and spent less time shoaling. Staying away from the group enabled them to establish and defend territories and increased foraging success, while also increasing predation risk.

Individual differences in social learning may be the result of risk-reward trade-offs associated with differences in foraging objectives. These differences emerge, for example, where parental investment is asymmetric or where reproduction is skewed [19, 20]. In species with low male parental investment or high reproductive skew, males often experience intense intrasexual competition as reproductive success depends on access to females [19]. Because male body condition often correlates with fighting ability and male-male contest success, males often engage in riskier foraging behaviour to maximise their energy intake and competitive ability [21, 22]. Male African elephants, for example, are more likely than females to engage in crop raiding [23]. Although risky, crop-raiding is associated with adult body size and mating success. Similarly, bachelor groups of male African buffalo accept increased predation risk in habitats with low visibility in return for higher energy gains [24].

On the other hand, females generally experience lower reproductive skew than males. Their reproductive success depends on successfully raising offspring. Therefore, reproductive output is limited by high energetic demands due to pregnancy, lactation, and parental care [25], as well as by the need to reduce the risk of predation and energetic shortfall, which could be fatal to their offspring. Females also strive to maximise energy input but prioritize low-risk strategies, even if that reduces potential gains. Preferences for risk-averse foraging strategies in females have been documented in several species. Female capuchins, for example, are more likely than males to be central in their group during foraging, where predation risk is lower [26]. A similar observation has been made in sticklebacks where gravid females were less likely to switch between food patches and spent more time in cover than non-gravid females or males did [13]. Also, female capuchins are more likely to feed on smaller, embedded, and more reliably found invertebrates, while males spend more time searching for and processing of large vertebrate prey [27, 28].

Aside from the risk directly associated with foraging, individuals have to vet different information sources to make foraging decisions and accomplish their foraging objectives. Where animals forage in groups they obtain information either individually (active searching) or socially (by observation of or interaction with others). Most resources are limited, and so individuals must compete for them. As social learning implies that at least one other individual has knowledge of the resource, social learners are more likely to be exposed to competition; this reduces the yield of social learners compared to individuals that rely less on social information [29]. However, copying others should also reduce the risk of exploiting resources with very low returns, and thus reduce the risk of energetic shortfalls [30, 31]. Whether individuals use social information should, therefore, depend on their risk-taking behaviour (or foraging objectives), where risk is determined by the amount and variance of expected returns [32, 33]. Thus, we expect that variation in foraging objectives due to risk sensitivity favours differences in social information use.

Surprisingly, despite the empirical evidence for individual differences in foraging objectives, how social learning evolution is affected by varying energetic requirements has not been studied [10, 34]. To investigate whether differences in the reproductive payoff derived from collected resources affect the optimal use of social information, we extended a previously published model of social learning to include reproductive skew [35]. We studied reproductive skew as this is a well-established source of differences in energetic requirements. We modelled sexes because these provide a case where individuals in the same population have different foraging objectives to maximise their fitness, i.e. either maximising resource return or minimising risk of energetic shortfall. However, sexes are not unique in causing differences in foraging objectives: we would expect more risk prone behaviour whenever reproductive output is more skewed. Finally, we framed our model in a foraging context where information refers to the quality of a resource patch, but the model would also apply if information described the expected return from different foraging techniques.

## 2 Model

In our model, individuals collect payoffs from patches in their environment. They acquire information about patches by individual or social learning, and they receive payoffs when they exploit those patches. Patches differ in payoffs, and individuals attempt to maximise their collection rate by exploiting the most profitable patches. Where multiple individuals exploit the same patch, payoffs are shared equally (i.e., exploitative scramble competition). We found the evolutionary equilibrium between individual and social learning by letting individuals die randomly or when reaching a certain age and replacing them through differential reproduction of adults as described below. Individuals must meet certain criteria in order to reproduce. By controlling those criteria, we controlled reproductive skew. Below we describe the model in more detail.

We simulated worlds with *N* individuals and *M* patches, where patch *m* provides a payoff *π*_*m*_. Payoffs are drawn from one of two different gamma distributions, which have the same mean expected payoff 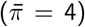, but different variances (*σ*^2^). We refer to environments with small variance in payoffs as even worlds (*σ*^2^ = 1) and those with large variance as uneven worlds (*σ*^2^ = 100).

Time is divided into rounds. Payoffs replenish after each round. To simulate environmental turnover the payoff in each patch changes with probability *τ* after any given round. In each round, each individual either learns (with probability 0.2) or exploits a patch (with probability 0.8). Individuals can only exploit resource patches they have previously learned about.

Individuals learn either individually (by sampling a random patch), or socially (by randomly sampling from the individuals that exploited patches in the previous round and learning about the patch exploited by one of those individuals). In any learning turn, individual *i* learns socially with probability *α*_*i*_. Thus, we assumed that individuals do not change their use of social learning in a context-dependent manner [8-10]. In the results presented in the body of the paper, individuals are either strict individual (*α* = 0) or strict social learners (*α* = 1), which allows us to compare foraging success between pure strategies. However, we also ran simulations where individuals probabilistically choose between individual and social learning (i.e., individuals can have *α*s between 0 and 1) and found the results to be qualitatively similar (see Appendix C in Supporting Information).

Through learning, individuals obtain information about the current payoff of a patch, which we refer to as the expected payoff *q*. Because individuals compete for resources, the expected payoff of a patch m is *q*_*m*_ = *π*_*m*_*/n*_*m*_, where *n*_*m*_ is the number of individuals currently exploiting the patch (where *n*_*m*_ = 0,*q*_*m*_ = *03C0*_*m*_). The expected payoff is stored in the individual’s memory. Information about a patch is retained until it is updated (that is, when an individual exploits that patch or learns about it again in a learning turn) or until the individual dies. An individual’s information about a patch may be incorrect if the patch value or occupancy has changed since the information was collected.

In each turn in which it exploits resources, an individual chooses the patch form its memory with the highest expected payoff. Exploiting patch *m* yields payoff *p*_*m*_ = *π*_*m*_*/n*_*m*_. We assumed that an individual’s ability to successfully reproduce depends on its resource collection rate, which is commonly used as a fitness proxy in theoretical models (see for example [10]). As recent payoffs are more relevant to current reproductive success than payoffs collected further in the past, we allowed reproductive success to depend on payoffs from the previous *δ* rounds. We weighted payoffs relative to how long ago they were collected using a weighted moving average. We defined the fitness proxy *F*_*i*_ of an individual *i* in round *t* of its life as:

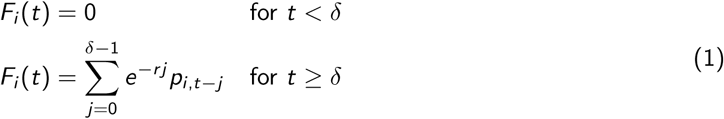

where *r* is a temporal discount rate. The first part of eq. (1) implies that individuals reach reproductive maturity at age *δ*. We ran simulations with window size *δ* = 5 and temporal discount rate *r* = 0.2 (figure 3,4).

Eq. (1) reduces variability in the fitness proxy by making it a weighted sum across rounds. This is biologically reasonable for systems in which individuals make frequent decisions about patches or foraging strategies and where body condition is a result of accumulated returns. However, to show that our results do not depend on this assumption, we analysed a simpler version of the model in which the fitness proxy in any round is simply the resources collected in that round (i.e., *F*_*i*_ (*t*) = *p*_*i,t*_, see figure S2). This is another common assumption [36-38] and is plausible, for example, if the decision individuals make is the choice of a seasonal foraging range and if each individual’s ability to raise offspring depends on the resources it collects in that season (see Appendix B).

Individuals die randomly (with probability 0.01 in each round) or deterministically (when they reach the maximum age of 100 rounds). Through reproduction each dead individual is replaced in the population, keeping the population size constant. The new individual inherits a social learning propensity *α* from its parent(s).

Next, we let the proportion of social learning evolve in systems with different levels of reproductive skew. Because the relative fitness effect of individual and social learning is frequency dependent, we can find the equilibrium proportion (at which both learning modes have the same fitness) by mimicking an evolutionary process. That is, reproduction is tied to foraging success. We achieved a skewed reproduction by ranking individuals according to their fitness proxy in the current round, where only individuals in top-ranked proportion *β* ∈ [0.01, 1] can reproduce. Decreasing *β* increases reproductive skew. All individuals in the top *β* proportion of the population are equally likely to reproduce. This might represent a case in which individuals must simply achieve a physiological threshold to reproduce (minimum income criterion). In some systems, individuals that face physiological constraints can reproduce whenever they collect enough resources to meet a minimum body condition threshold for breeding [39-41]. In Appendix B, we show that our results hold when the probability that an individual in the top *β* proportion of the population reproduced is proportional to its fitness proxy (relative income criterion). This might represent situations when reproduction is socially constrained and requires successful competition for partners, and the most competitive or high ranking individuals with good body conditions are most able to reproduce [42, 43].

In simulations with evolving social learning frequency, reproduction is either asexual or sexual. In the case of asexual reproduction all individuals of a population experience the same reproductive constraints. Here, each offspring directly inherits its social learning propensity *α* from its parent. To prevent stochastic extinction of learning modes, mutation from *α* = 1 to *α* = 0 or vice versa occurs with probability *μ* = 1*/N*.

Where reproduction is sexual, individuals within the same population experience different reproductive constraints. For simplicity, we refer to the subset of individuals experiencing higher reproductive skew as ‘males’ and the subset experiencing lower reproductive skew as ‘females’. However, in nature those labels might be reversed or tied to characteristics other than sex. Under this formulation of the model, individuals can reproduce whenever they are not in the bottom 1 - *β* proportion of the foragers of their sex. Thus, the minimum resource requirement for reproduction for each individual is calculated relative to the foraging success of other individuals of its sex. In nature, individuals may also need to achieve some fixed minimum resource collection in order to reproduce. In Appendix B we show that results are qualitatively similar when this is true.

In simulations with sexual reproduction, individuals are either male or female and their social learning propensity is under control of a diploid genome. The social learning propensities in males and in females are each controlled by a single locus, and the loci that control male and female social learning propensities are on different chromosomes (i.e., they assort independently). Thus, the social learning propensities of each sex can evolve independently, and there is no potential for sexual conflict. If an individual has different alleles at the loci that govern the social learning propensity for its sex, its social learning propensity is the mean of those alleles.

To replace a dead individual with a new individual one male and one female are randomly drawn from the top *β_m_* proportion of males and the top *β_f_* proportion of females to be the parents. The same individual can be chosen as a parent multiple times in the same round. Next, a sex is assigned to the offspring with equal probabilities to be male or female. The offspring receives one randomly chosen allele from each parent at each locus, with free recombination. As in models with asexual reproduction, we simulated cases in which alleles are constrained to have values of *α* = 0 or *α* = 1 (see Appendix C for cases in which alleles can take any value from 0 to 1). Mutation occurs independently for each inherited allele with probability *μ* = 1*/N*.

### Model Iterations

We first simulated populations with fixed social learning frequencies and tested whether social learning is a risk-averse strategy by asking whether it yields less variable returns than individual learning (figure 1, 2). In these simulations, individuals that died were replaced by copies of themselves with empty memories and no collected resources. Thus, there was no reproductive skew or selection. As expected, we found that the returns of social learners were less variable than those of individual learners, because social learners were less likely to exploit poor quality patches. We then introduced reproductive skew in asexually reproducing populations, where reproductive skew was the same for all individuals. This provides a baseline for the effect of reproductive skew on social learning evolution (figure 3,4). Finally, we modelled worlds with sexual reproduction between individuals who differ in reproductive constraints. This allowed us to study the evolution of social learning when individuals with different reproductive constraints are present in the same population, and how the distribution of social learning in the population changes when differences in reproductive skew are high or low.

**Figure 1:**
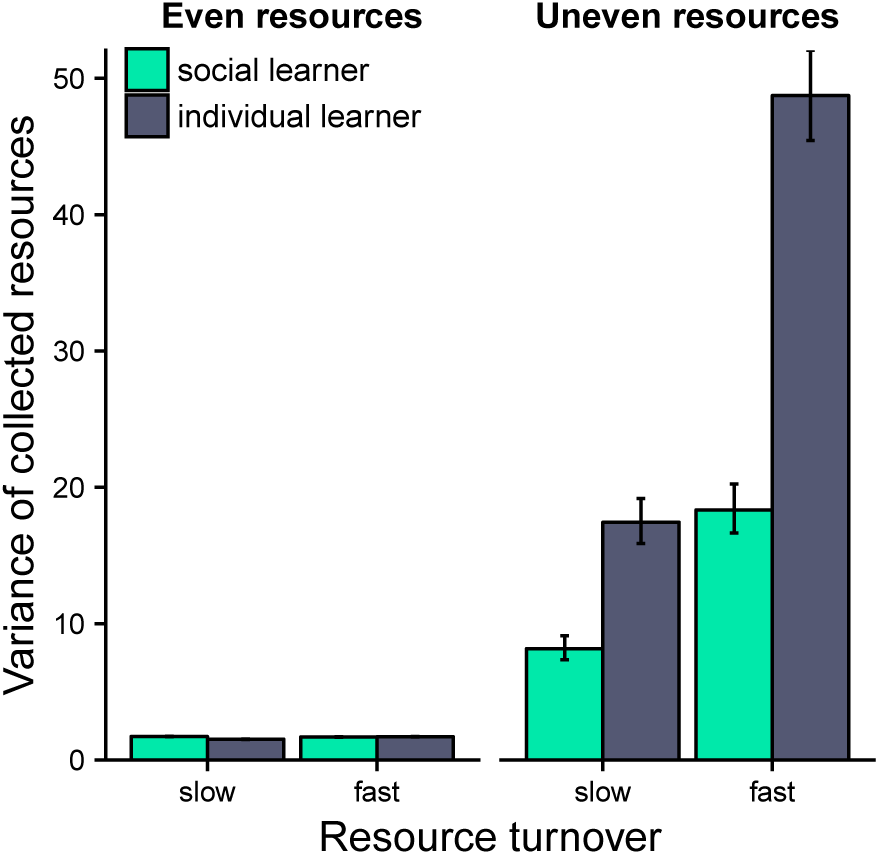
In simulations where the proportion of individual to social learners remained fixed at 50%, individual learners had higher variability in collected resources when resources were unevenly distributed. When resources were evenly distributed, variability was similar for the learning strategies. Data shown are collected resources of all exploiting individuals in the last simulation round (i.e., 5000). Resource turnover rates (slow: *τ* = 10^-2^, fast: *τ* = 10^-1^, resource distributions (even: *σ*^2^ = 1, uneven: *σ*^2^ = 100). Error bars indicate 95% confidence intervals.

**Figure 2:**
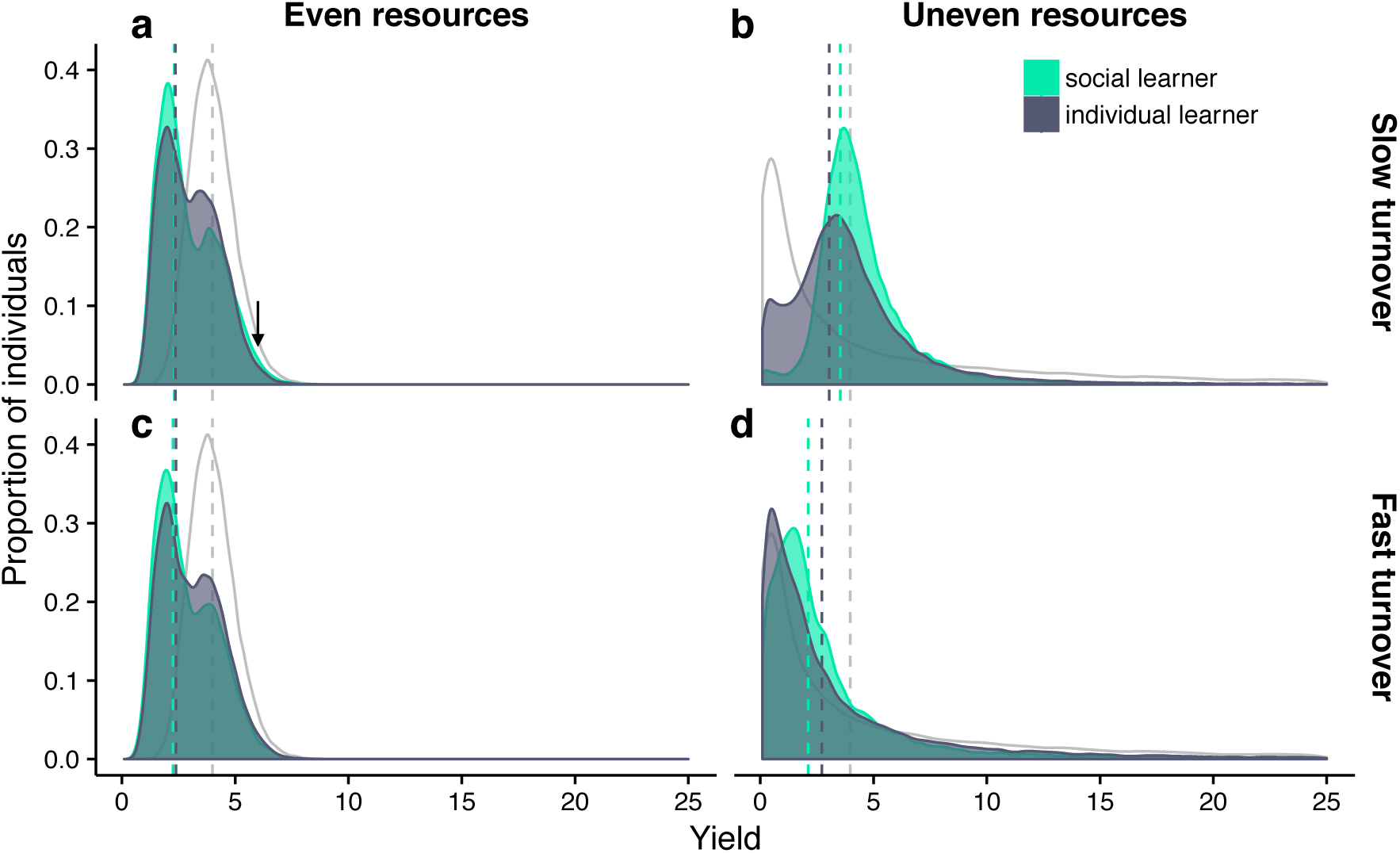
Distribution of resources collected per round by social and individual learners under different resource distributions (even: *σ*^2^ = 1, uneven: *σ*^2^ = 100), and turnover rates (slow: *τ* = 10^-2^, fast: *τ* = 10^-1^). Where resources are evenly distributed (a,c), most individuals either share a patch with one other individual (peak at 2) or are alone in a patch (peak at 4). A small proportion of social learners collects more resources than individual learners where resources are slowly turning over and are evenly distributed (arrow in a). For unevenly distributed resources (b,d), there are more individual learners than social learners collecting resources from poor patches, as well as from highly profitable patches. Data are truncated along x-axis at 25; grey line indicates environmental resource distribution; dashed vertical lines indicate distribution means.

**Figure 3:**
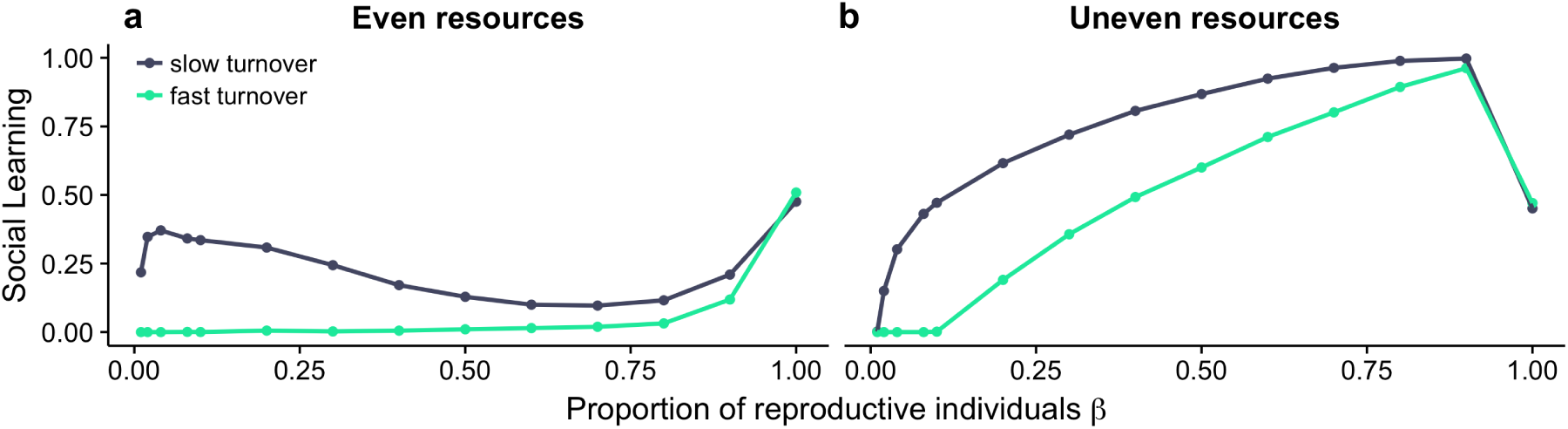
Evolving proportions of social learning for different resource distributions (even: *σ*^2^ = 1, uneven: *σ*^2^ = 100) and environmental turnover (slow: *τ* = 10^-2^, fast: *τ* = 10^-1^) in the minimum income scenario. Where reproductive skew is highest (small values of *β*) individuals rely mostly on individual learning, with the exception of slowly changing, evenly distributed resources. At low reproductive skew (large values of *β*) there is more social learning. In addition to reproductive skew, the proportion of social learning also depends on the resource turnover and the resource distribution. There is more social learning when resources are unevenly distributed and environmental turnover is slow. Results are qualitatively similar in the relative income scenario (figure S3).

**Figure 4:**
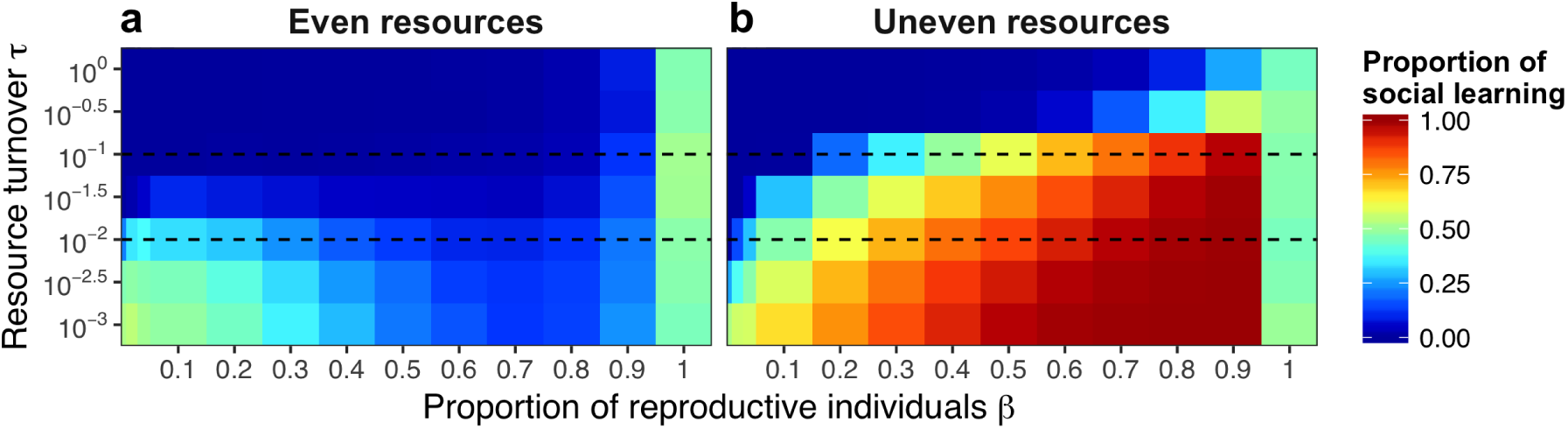
Environmental predictability and reproductive skew affect social learning under different resource distributions. Where resources are unevenly distributed (b) reliance on social learning increases as reproductive skew and environmental turnover decrease. Where resources are evenly distributed and frequently changing (a) individuals almost exclusively rely on individual learning. However, for very slow environmental turnover social learning can evolve even at high reproductive skew (see text for details). Dashed lines mark high and low resource turnover rates shown in figure 3. Results are qualitatively similar in the relative income scenario (figure S4).

### Model initialisation

For simulations with fixed proportions of social learning, we initialized worlds with 1000 patches and with 500 individual and 500 social learners for all combinations of *β* ∈ {0.01, 0.02, 0.04, 0.08, 0.1, 0.2, 0.3, …, 0.9, 1}, resource turnover *τ* ∈ {10^0^, 10^-0^.^5^, 10^-1^, …, 10^-3^} and resource variance (*σ*^2^ ∈ {1, 100}).

For simulations with sexual reproduction, populations were initiated with 500 females and 500 males. Zeros and ones were randomly assigned to the alleles that determine social learning propensity in both sexes. We ran simulations with sexual reproduction for intermediate turnover *τ* = 10^-1^.^5^, and high resource variance *σ*^2^ = 100, as these parameters support intermediate adoption of social learning in a population (see [35]).

All simulations ran until either all individuals shared the same value for *α* or a maximum of 5,000 rounds was reached. Each parameter set was repeated 200 times.

### Code availability

All work presented here was done in R [44]. Software source code for the computer model and for the accompanying figures will be made freely accessible at an online repository upon acceptance.

## 3 Results

### Social learning is a risk-averse strategy

In our first analysis, we kept the proportion of social and individual learners fixed at 50% to compare how the variance in collected resources differed between individual and social learners (figure 1). When resources were evenly distributed, variance in collected resources was low and did not differ meaningfully between social and individual learners (ratio of variance 1.04 and 0.94 for slow and fast turnover). When resources were unevenly distributed the variance in collected resources was higher among individual learners than among social learners for both turnover rates (ratio of variance 0.34 and 0.36 for slow and fast turnover). As variance in collected resources is a common measure of risk sensitivity, our results suggest that social learning is a risk-averse strategy, whereas individual learning is a risk-prone strategy.

To get a more detailed look at the distributions that underlie this pattern we plotted the collected resources for social and individual learners for the different environments in figure 2. When resources were unevenly distributed, the position of the distribution peaks in figure 2b,d show that there are on average more social learners foraging in more profitable patches, whereas the fatter tails of the individual learner distributions indicate that more individual learners foraged in the highest profit patches. The mean return rate was higher for social learning when resources turned over slowly, and for individual learning when resources turned over more quickly (compare vertical lines in figure 2b,d).

When resources were evenly distributed, resource collection followed a bimodal distribution with one peak for individuals that are alone in a patch and a second for those sharing a patch with another individual (figure 2a,c). For both turnover rates, there were more individual learners alone in patches compared to social learners, but the average amount of collected resources (vertical lines) was similar for the two learning types.

Surprisingly, at slow environmental turnover, there was a small proportion of social learners that foraged in slightly more profitable patches than individual learners (arrow in figure 2a). The explanation for why there are some social learners that are more successful in this situation is that when resources change slowly individuals approach an ideal free distribution. When resources are evenly distributed, and the population density is low, most patches have one individual. Consider a social learner that moves into a patch with another individual. If the individual in that patch knows of a better patch, it will leave, and the social learner is left alone. Otherwise both stay. If the social learner is the next to learn, it will move only if it finds a better patch to share. The social learner will continue moving to better and better patches until its co-occupant either leaves or dies. As it moves from patch to patch, it will eventually occupy a patch that is better than the average patch occupied by individual learners. A new individual learner benefits from the same process, but social learners are more likely to join another individual than individual learners. Eventually, an individual learner will find and accept a slightly poorer unoccupied patch, which is still better than sharing. Essentially, social learners drive individual learners out of, and therefore end up alone in, high-reward patches.

### Reproductive skew can affect the preference for social learning

The previous section shows that individual learning can increase maximum collected resources (figure 2), whereas social learning decreases income variability (figure 1). As reproductive skew rises in our model, reproductive success increasingly relies on achieving the highest resource collection. Therefore, we expect that individual learning will be favoured where reproduction is highly skewed. In contrast, where reproductive skew is low, individuals with less variable income will be favoured as they will more often have sufficient resources to reproduce. Therefore, we expect that social learning will be favoured where reproductive skew is low. Figure 3 and 4 show the results of simulations where all individuals in a population experience the same level of reproductive skew, *β*, for different values of *β* and different environmental conditions. We find that the amount of social learning is affected by reproductive skew and modulated by environmental turnover and resource distribution.

When resources were unevenly distributed, social learning evolved when reproductive skew was low, but individual learning evolved when reproductive skew was high (figure 3b). High-quality patches can be shared and still yield resources well above the lower part of resource collection distribution in the population, which favours social learning when reproductive skew is low. However, if reproductive skew is high, only the most successful foragers reproduce, which tend to be individual learners foraging alone in high-quality patches. Thus, high reproductive skew favours individual learning in uneven environments.

When resources were evenly distributed, individual learning was more likely to evolve (figure 3a). In this case, patch resources are close to the environmental mean and sharing a patch seldom provides enough resources for reproduction. Therefore, individuals gain more by foraging alone. Because individual learners are more likely to be alone in patches (figure 2), social learning remains at low levels.

As expected from previous work [35], individual learning dominated in quickly changing environments, whereas social learning dominated when environments changed slowly, especially if reproductive skew was low and resources were unevenly distributed (figure 4b). For slow environmental turnovers and even resources, social learning can evolve even at moderate to high reproductive skew (figure 3a, 4a). This unexpected outcome from our model is due to the small number of social learners that come to monopolise high quality patches as described for figure 2a. The result only holds for low population density where individuals are often alone in patches and disappears at higher population density (see Appendix A, figure S1).

With the exception of the limiting case without reproductive skew (*β* = 1) at which there is no selection and on average 50% of social learning evolves (due to symmetrical mutation), we find that for the majority of the parameter space uneven environments favour social learning more than even environments (compare figure 4a and 4b).

### Sexual reproduction increases learning differences

Our results suggest that populations with lower reproductive skew should use more social learning than populations with higher reproductive skew. To test whether this result holds when individuals within a population experience different levels of reproductive skew, we moved to the sexual version of our model where we divided the population into males and females (figure 5). We found social learning levels were low when reproductive skew was high (lower left of figure 5a-c), and high when reproductive skew was low for both males and females (upper right of figure 5a-c), which corresponds to the results of the asexual model. At intermediate levels of reproductive skew, we found that the competition from each sex influences the evolved social learning propensity in the other. Over most parts of the parameter space, females evolve to learn almost entirely socially, and males evolve to learn almost entirely individually. As reproductive skew decreases social learning increases faster in females than in males (compare the change in social learning along the main diagonal for males in figure 5a with that for females in figure 5b). As in the asexual simulations, we found that social learning evolves on average to a level of 50% in females for the limiting case *β_f_* = 1 (figure 5b), in which there is symmetrical mutation and no selection.

**Figure 5:**
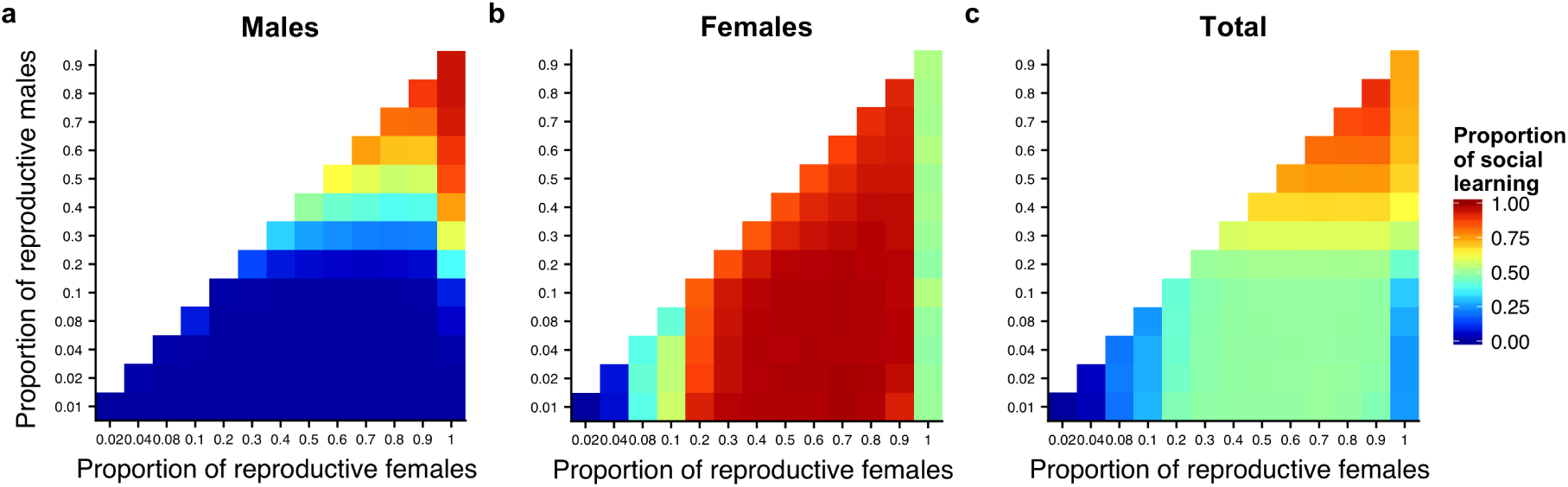
At high reproductive skew (bottom left of each panel), both males and females rely almost exclusively on individual learning, whereas at low reproductive skew (top left of each panel) social learning dominates (c). However, for intermediate levels of reproductive skew, we find that each sex adopts a more extreme strategy due to the presence of the other sex, often approaching either pure individual learning (males, a) or pure social learning (females, b). Results shown are from simulations where males have a higher reproductive skew (along the y axis, increasing values indicate more males being eligible to reproduce, and thus reproductive skew is smaller), than females (x-axis). The simulated environments had high patch payoff variance (*σ*^2^ = 100) and intermediate resource turn-over (*τ* = 10^-1.5^).

Overall, our results suggest that when both sexes share the same environment and differ in reproductive skew, differences in the use of social learning can become pronounced (figure 6).

**Figure 6:**
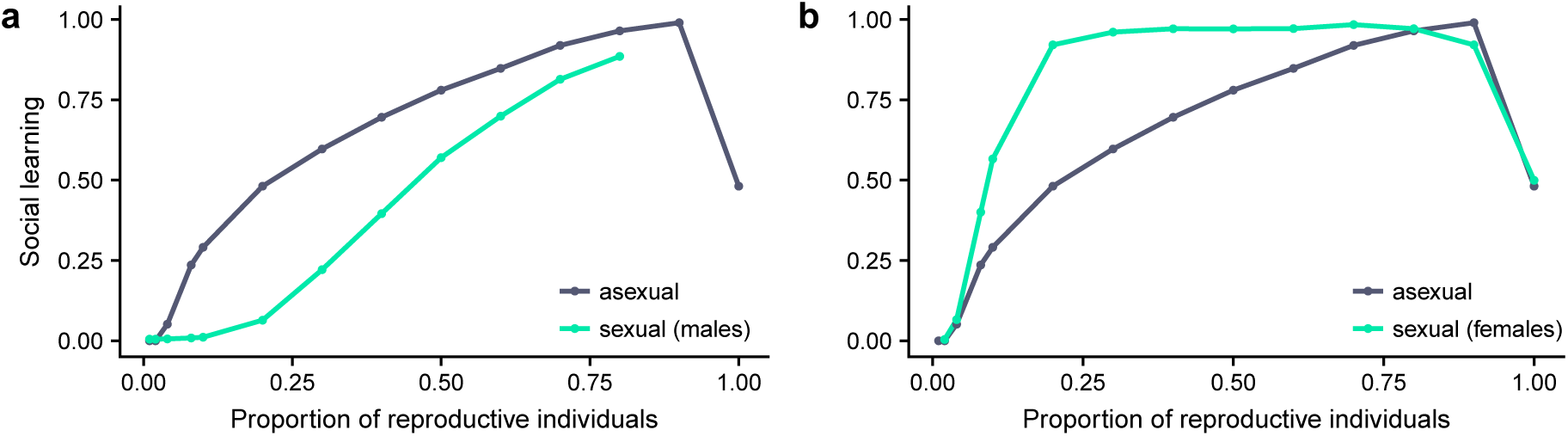
Where both sexes share the same resources males use less and females more social learning than would be expected in asexually reproducing populations. Results for sexuals are taken from simulations as shown in figure 5. Results for asexual simulations are identical to those in figure 4b, where *τ* =10^-1.5^, for males (a) to those in figure 5a, where *β_f_* = 0.9, and for females (b) to those in figure 5b, where *β_m_* = 0.01.

## 4 Discussion

We used an agent-based foraging model to investigate how reproductive skew affects the optimal use of social learning. We found that individual learning is a risk-prone and social learning a risk-averse foraging strategy, low reproductive skew favours social learning in the population, and differences in reproductive skew between the sexes can promote differences in the use of social learning. These results are moderated by the environment, as social learning is more advantageous when resources are unevenly distributed and temporarily predictable.

Foraging returns from individual learning were generally more variable than from social learning because individual learners sample patches across the environment and are more likely to encounter patches with extreme (high or low) amounts of resources compared to social learners. Social learners, in contrast, experience lower variance because they only copy information that was retained by other individuals. Although encountering a poor patch is less likely for social learners, social learners are also less likely to achieve very high returns as they are more often in occupied patches sharing resources with others. Caraco et al. [45] have shown that foraging socially decreases food-discovery variance. Similarly, a study on European starlings (*Sturnus vulgaris*) reported that information scrounging was negatively related to food-intake variance [31]. This suggests that, in nature, social learning is a risk-averse strategy with more reliable but on average lower returns, whereas individual learning is a risk-prone strategy with less predictable but potentially higher returns, and is consistent with a previous theoretical model on social learning evolution [30].

How individuals respond to risk should depend on their foraging objectives, which depend in turn on their reproductive strategy. Individuals that have much to gain from improving their condition and little to lose if their condition deteriorates should use risk-prone strategies, whereas those that have little to gain from an improved condition and much to lose from deterioration should be risk-averse [46-48]. Previous studies have shown that individuals can alter their foraging to match their life histories, in that individuals that require more resources to successfully reproduce increase foraging returns by choosing more risk-prone foraging strategies [21, 23, 24, 46, 49]. Our work advances this understanding by tying it to individual and social learning. Where reproductive skew was high individuals relied almost exclusively on risk-prone individual learning. Conversely, at low reproductive skew individuals relied more on social learning when resources were unevenly distributed and stable over time. In this case, social learners are more likely than individual learners to collect sufficient resources for reproduction (i.e., be in the *β* percentile). This conforms to earlier findings that individuals are generally risk-averse if expected returns exceed required returns [48, 50].

Consistent with previous work, there was more social learning when resources were unevenly distributed [30, 35, 51, 52]. When resources were more evenly spread among patches, social learning was less adaptive, as sharing a patch in this environment reduces the expected return below the environmental average. A surprising outcome of the model is that for very stable and evenly distributed resources, we found social learning at medium to high levels of reproductive skew, as long as population density was not too high. In such cases, the best foragers were social learners, but these best foragers were only slightly better than their competitors. In nature, this pattern would favour social learning only if fitness is very sensitive to small differences in foraging success. This may be true in at least some systems. For example, three-spined stickleback females (*Gasterosteus aculeatus*) choose mates based on both visual and olfactory cues. However, when the water is very clear, females rely almost exclusively on visual cues [53], and even small differences in diet-dependent colouration could strongly affect male reproductive success.

Here, we provide a theoretical framework suggesting that differences in social learning can be explained by differences in reproductive skew and the resulting differences in risk-taking. These differences have been documented both in humans and non-human animals, usually with higher risk aversion in females than in males [54-62]. However, we do not wish to suggest that our results imply general differences in social learning efficacy or capacity between males and females. Instead, our model was motivated by the observation that individuals differ in their foraging objectives to accommodate their energetic requirements. Similarly, these results could apply at the species level, depending on reproductive skew and the distribution and turnover of resources. These objectives might be consistently different among individuals but can also change over time and between contexts. For example, an individual could be more receptive for or attentive to certain sources of information depending on its ecological or life-history condition. This reflects the case in the previously mentioned example with sticklebacks, where reproductively active females used social learning more often, while there was no difference between non-reproductive females and males [13].

Previous theoretical work commonly assumes that individuals maximise resource intake, and that the same increase in payoffs translates to the same potential increase in fitness for all individuals [9, 10]. However, we have shown that our understanding of social learning evolution benefits from explicitly modelling individual differences in energetic requirements, for example, by including reproductive skew. This allowed us to observe that in populations with sexual reproduction males and females become more extreme in their learning strategies, an effect that is absent in previous models. Based on our results, sex differences in socially acquired foraging and tool-use skills that have been reported in bottle-nose dolphins [63] and chimpanzees [64, 65] might be better explained by differences in foraging objectives than by general learning differences. In chimpanzees, for example, females might be more receptive to foraging behaviours because their reproductive success relies on obtaining enough food to bear and raise young, whereas males might be more attentive to dominance behaviours and forming alliances to establish their position in the group [65]. Similarly, in bottle-nose dolphins females might be more likely to acquire foraging behaviours that are locally adaptive, whereas males ensure mating opportunities by forming alliances with other males which require them to roam outside of areas where sponge-carrying would be adaptive [66, 67].

One application of our results is the hypothesis that males and females may contribute differently to the emergence, spread, and accumulation of certain cultural traits. For example, in some primates, innovation (a form of individual learning) is found more often in males than in females [68], whereas females appear to use more social learning in chimpanzees [64, 69, 70], bonobos [71], and macaques [72]. Lind & Lindenfors [70] note that the number of cultural traits in chimp communities correlated with the number of females but not males. The authors argue that this is likely because females transferred between communities and brought novel socially learned traits into communities and refer to female chimps as *cultural carriers*. Whether this extends to human cultural evolution in the past is unclear. While studies involving modern humans report sex differences in social information use [62, 73], studies on social learning efficacy found no sex differences [74, 75]. Even if sex differences in social learning existed in our ancestors, these may have disappeared through selection as cultural complexity increased. In a society with highly complex or large numbers of cultural traits, successful social learning would affect survival and ultimately reproductive success. More ecologically relevant studies might give us an opportunity to find differences in social learning where they exist [76]. Future work should consider the implications of sex differences in risk-taking on learning strategies, and how they could lead to a new understanding of sex roles in the evolution of culture.

**Table 1:**
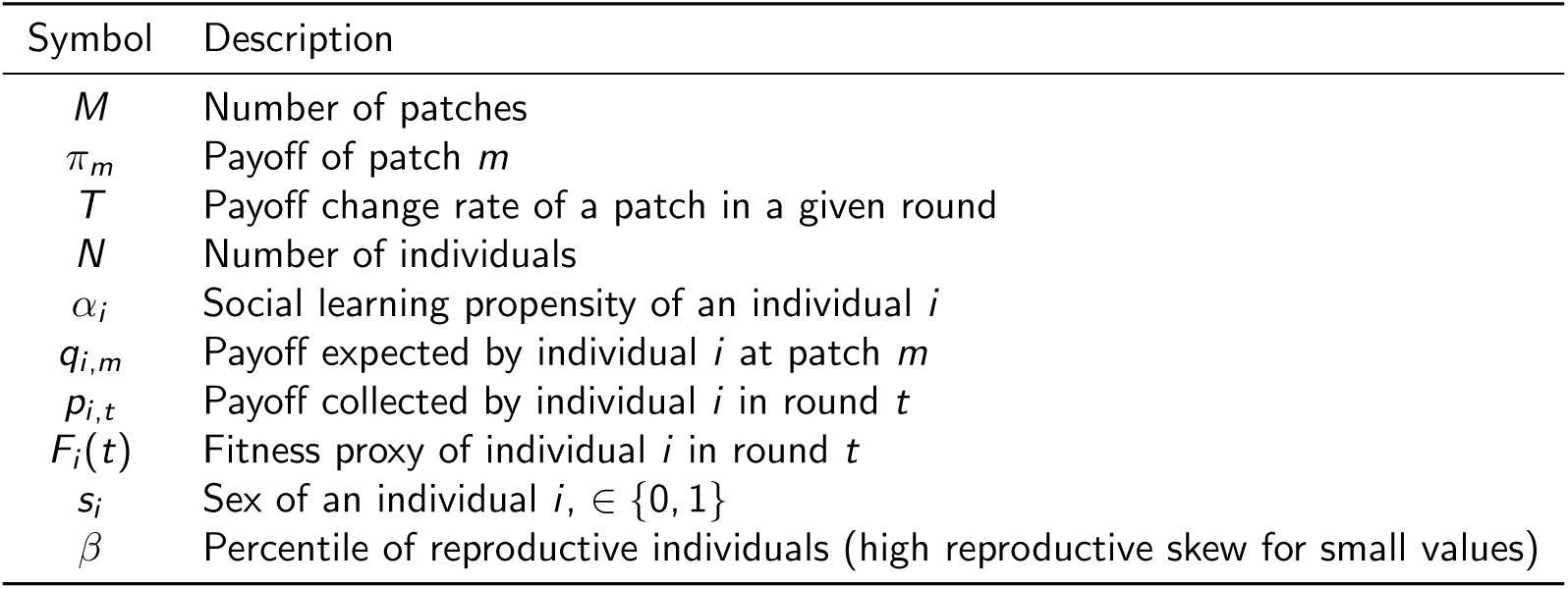
Symbols and definitions used in the model.

| Symbol | Description |
| --- | --- |
| $M$ | Number of patches |
| $\pi_m$ | Payoff of patch $m$ |
| $T$ | Payoff change rate of a patch in a given round |
| $N$ | Number of individuals |
| $\alpha_i$ | Social learning propensity of an individual $i$ |
| $q_{i,m}$ | Payoff expected by individual $i$ at patch $m$ |
| $p_{i,t}$ | Payoff collected by individual $i$ in round $t$ |
| $F_i(t)$ | Fitness proxy of individual $i$ in round $t$ |
| $s_i$ | Sex of an individual $i$, $\in \{0, 1\}$ |
| $\beta$ | Percentile of reproductive individuals (high reproductive skew for small values) |

## Acknowledgement

MS was supported by a Royal Society Research Grant and in part by a grant to Erol 630 Akçay from the Army Research Office (W911NF-12-R-0012-03), SS is supported by a Royal Society University Research Fellowship.

